# Transient Nodal signalling in left precursors coordinates opposed asymmetries shaping the heart loop

**DOI:** 10.1101/854463

**Authors:** Audrey Desgrange, Jean-François Le Garrec, Ségolène Bernheim, Tobias Holm Bønnelykke, Sigolène M. Meilhac

## Abstract

The secreted factor Nodal has been shown to be a major left determinant. Although it is associated with severe congenital heart defects, its role in heart morphogenesis has remained poorly understood. Here, we report that Nodal is transiently active in precursors of the mouse heart tube poles, before the morphological changes of heart looping. In conditional mutants, we show that Nodal is not required to initiate asymmetric morphogenesis. We provide evidence of a heart-specific random generator of asymmetry that is independent of Nodal. Using 3D quantifications and simulations, we demonstrate that Nodal functions as a bias of this mechanism: it is required to amplify and coordinate opposed left-right asymmetries at the heart tube poles, thus generating a robust helical shape. We identify downstream effectors of Nodal signalling, regulating asymmetries in cell proliferation, cell differentiation and extra-cellular matrix composition. Our work provides novel insight into how Nodal regulates asymmetric organogenesis.

## Introduction

Left-right asymmetric organogenesis is essential for vertebrates. Impairment of left-right patterning leads to human diseases such as the heterotaxy syndrome, affecting the asymmetry or concordance of the positions of visceral organs (Van Praagh, 2006). Heterotaxy is associated with complex congenital heart defects which determine the prognosis of patients (Lin et al., 2014). How the bilateral symmetry is broken in the early embryo is now well established. This involves a left-right organiser, referred to as the node in the mouse, which forms as a pit of ciliated cells at E7.5 (see Lee and Anderson, 2008; Shiratori and Hamada, 2006). The motility of cilia generates a leftward fluid flow (Nonaka et al., 1998), which is required between the 1 to 6 somite stages for the asymmetric expression of components of Nodal signalling. *Dand5*, encoding a Nodal antagonist, is the first gene asymmetrically expressed, on the right side of the perinodal region (Kawasumi et al., 2011; Marques et al., 2004). As a result, Nodal, a secreted factor of the TGFβ family, becomes active only on the left side of the perinodal region, as detected by the phosphorylation of the Smad2 transcription factor (Kawasumi et al., 2011). It stimulates its own asymmetric expression in the left perinodal region and the left lateral plate mesoderm (Brennan et al., 2002; Saijoh et al., 2003). *Nodal* expression is initiated lateral to the node, then propagates anteriorly and posteriorly in the left lateral plate mesoderm by auto-activation (Lowe et al., 1996; Vincent et al., 2004). Nodal also activates its own antagonists *Lefty1/2*. This negative feedback loop ensures transient expression of *Nodal* in the lateral plate mesoderm, where heart precursors are localised, between the 3 and 6 somite stages. *Nodal* is not expressed within the heart tube (Collignon et al., 1996; Vincent et al., 2004). Deletion of the asymmetric enhancer (ASE) of *Nodal* dramatically reduces left-sided *Nodal* expression and impairs the formation of the heart and lungs (Norris et al., 2002). Similarly, the conditional inactivation of *Nodal* in the mesoderm impairs left-right asymmetry of the heart, lungs, spleen and stomach (Kumar et al., 2008). Nodal is thus a major left determinant, required in the lateral plate mesoderm for asymmetric organogenesis. The phenotype of *Nodal* mutants is either symmetrical, i.e. right isomerism as seen in the lungs, spleen and atria, or randomly lateralised, as seen in the stomach, gut, heart apex or heart tube (Brennan et al., 2002; Kumar et al., 2008; Lowe et al., 2001; Saijoh et al., 2003). Beyond its asymmetric expression in cell precursors, it has remained unclear how Nodal regulates morphogenesis.

Asymmetric organogenesis has been proposed by Brown and Wolpert (1990) to occur in two steps: (1) a symmetry-breaking event, transforming molecular chirality into a left-right bias, which coordinates laterality throughout the embryo ; (2) organ-specific random generators of asymmetry, which are modulated by the left/right bias, to generate asymmetric organ shapes. Nodal signalling provides such a left-right bias, whereas random generators of asymmetry are still poorly characterised. The primordium of the heart is a tube, which grows by addition of precursor cells from the heart field to the arterial (cranial) and venous (caudal) poles (Domínguez et al., 2012; Kelly et al., 2001). The heart becomes asymmetrical during the process of heart looping, which transforms the tubular primordium into a helix at E8.5. This is crucial to position the cardiac chambers relative to each other and thus establish the double blood circulation (see Desgrange et al., 2018). From 3D (dimensions) reconstructions in the mouse embryo, we have previously characterised the spatiotemporal dynamics of heart looping, and established specific staging criteria. A model for a heart-specific random generator of asymmetry was proposed (Le Garrec et al., 2017). We showed that the heart tube grows between fixed poles, which is compatible with a buckling mechanism, able to generate random deformations. We also uncovered sequential and opposed asymmetries at the poles, which can bias the buckling to generate a helix: a rightward rotation of the arterial pole at E8.5f, followed by an asymmetric ingression of heart precursors at the venous pole at E8.5g. Finally, the progressive breakdown of the dorsal mesocardium provides an additional mechanical constraint that reinforces looping. On these bases, we have generated a computer model, which can predict the shape of the heart loop from a combination of values of the morphogenetic parameters. However, the molecular mechanisms of the left-right asymmetries at the heart tube poles remain unknown.

Nodal (Brennan et al., 2002; Kumar et al., 2008; Lowe et al., 2001; Saijoh et al., 2003), or components of Nodal signalling such as the transcription factor Foxh1, the co-receptor Cfc1, or the protease Furin (Roebroek et al., 1998; von Both et al., 2004; Yan et al., 1999), have been shown to control the direction of the heart loop. These studies have focused on the binary looping direction, as a readout of the symmetry-breaking event, but have not addressed the specific shape of the heart helix, as a readout of the heart-specific generator of asymmetry. In addition to heart looping, left-right patterning of cardiac precursors is important for the left/right identity of atrial chambers and the morphogenesis of the outflow tract. Clonal analyses of myocardial cells have shown a differential origin of the pulmonary trunk and aorta from left and right precursors respectively (Lescroart et al., 2010; 2012). In keeping with the spiralling of the great arteries in the mature heart, the outflow tract undergoes a rightward rotation (Bajolle et al., 2006), which follows that of the arterial pole during heart looping (Le Garrec et al., 2017). Regionalisation of the outflow tract prefigures the separation of the great arteries, as marked by *Sema3c* (Théveniau-Ruissy et al., 2008). Outflow tract defects have been associated with *Nodal* mutations in mouse models (Kumar et al., 2008; Lowe et al., 2001; Saijoh et al., 2003) and in patients, including a higher occurrence of pulmonary atresia and transposition of the great arteries (Bouvagnet and de Bellaing, 2016; Mohapatra et al., 2008).

A pending question is the identification of the molecular effectors downstream of Nodal signalling, and how they control asymmetric cell behaviour during organogenesis. Targets of the Nodal pathway have been mainly identified in the context of zebrafish gastrulation (Bennett et al., 2007), ES cell differentiation (Brown et al., 2011) or in cell cultures (Coda et al., 2017; Guzman-Ayala et al., 2009), but not in the context of left-right asymmetric organogenesis. A target gene of Nodal signalling in the left lateral plate mesoderm is the isoform *Pitx2c*, which is expressed asymmetrically in the heart tube, mainly in the inner curvature (Campione et al., 2001; Furtado et al., 2011). However, *Pitx2* or its isoform *Pitx2c* are not required for the proper direction of heart looping (Liu et al., 2002; M. F. Lu et al., 1999), highlighting the existence of other Nodal target genes during this process. Nodal is not always required for asymmetric organogenesis, as shown in the fly (see Coutelis et al., 2014; Spéder et al., 2006). Other determinants than Nodal have been identified, for example on the right side (Ocaña et al., 2017; Sivakumar et al., 2018) or acting in parallel (Pai et al., 2017).

Here, we address the specific role of Nodal for mouse heart looping with a high spatiotemporal resolution. We mapped the cardiac precursor cells expressing *Nodal* and determined the time window of Nodal activity. We generated conditional *Nodal* mutants, using *Hoxb1^Cre^*to target the mesoderm. We combined quantifications of the heart shape in 3D and computer simulations to assess how Nodal controls the heart-specific generator of asymmetry. Our results show that the generator of asymmetry is independent of Nodal, and that Nodal is required to amplify and bias pre-existing asymmetries at the tube poles. We show that *Pitx2c* is not required for mediating the biasing signal of Nodal. By transcriptomic analyses, we identified targets of Nodal, involved in the regulation of asymmetric cell behaviour at the heart tube poles. Thus, Nodal is essential for the robust rightward looping of the heart, which underlies the alignment of cardiac chambers.

## Results

### *Nodal* is expressed in myocardial precursors contributing to a quarter of the heart poles

Since *Nodal* is transiently expressed in the left lateral plate mesoderm and not within the heart tube, we investigated whether *Nodal*-expressing cells are genuine heart precursors. We took advantage of the *Nodal-ASE-lacZ* transgenic line, in which the β-galactosidase is produced under the control of the asymmetric enhancer of *Nodal*. The known perdurance of *lacZ* mRNA and/or β-galactosidase (Echelard et al., 1994) provides a pulse-chase readout of *Nodal* expression in the left lateral plate mesoderm. By 3D imaging of stained embryos at progressive stages of heart looping, β-galactosidase positive cells were found not only in the left heart field, but also at the heart tube poles : in the left sinus venosus, the myocardium of the dorsal left atrium, of the superior atrio-ventricular canal and of the left outflow tract (Fig. 1A-F, Movie S1). A sharp boundary was observed within the right ventricle and at the entrance of the left ventricle, such that the two ventricles were largely devoid of β-galactosidase staining (Fig. 1G). In the outflow tract, the domain positive for *Sema3c,* a marker of the pulmonary trunk (Théveniau-Ruissy et al., 2008), is broader and includes that of *Nodal-ASE-lacZ* (Fig. 1H-I). After segmentation of the heart tube myocardium, we measured that a fraction of 17% ±5 (n=16) is colonised by β-galactosidase positive cells, at all stages between E8.5e and E9.5 (Fig. 1J). If we consider only the heart regions colonised by β-galactosidase positive cells, i.e. discarding the ventricles, the fraction increases to 26% ±4 (n=4 at E8.5j and E9.5). Thus we show that left cardiac precursors, which contribute to the ventricles, do not express *Nodal*, whereas left cardiac precursors expressing *Nodal* contribute to a quarter of the cells in the heart tube poles.

**Figure 1.**
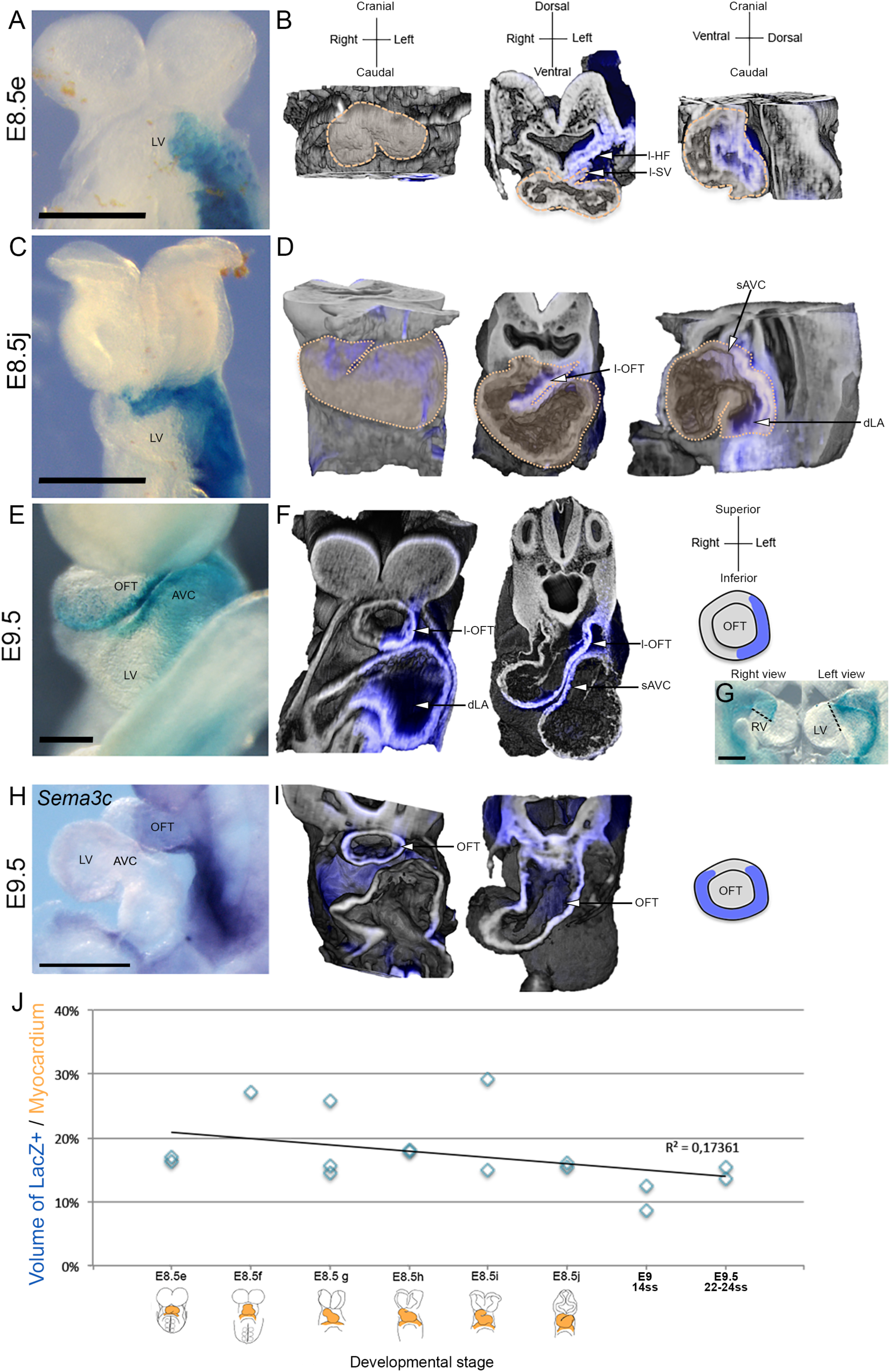
Tracing cells that have expressed Nodal with the *Nodal-ASE-lacZ* transgene. (A-G) Brightfield images (A, C, E, G) and 3D images by HREM (B, D, F) of *Nodal-ASE-LacZ* embryos at E8.5e (A-B), E8.5j (C-D) and E9.5 (E-G). Ventral views (left panels), transversal (third panel) and sagittal (right panel) sections are shown, with β-galactosidase staining in blue. The orange dotted line outlines the heart tube. Regionalisation of the staining in the outflow tract is schematised on the right (F). The black dotted lines (G) highlight sharp boundaries between β-galactosidase-positive and negative regions. See also Movie S1. (H-I) Brightfield image (H) and 3D images by HREM (I) of *Sema3c in situ* hybridisation (in blue). (J) Quantification of the β-galactosidase-positive myocardium volume at the indicated stages of heart looping. The low correlation coefficient (R^2^) indicates a constant fraction at the different stages. Scale bars: 200µm. dLA, dorsal left atrium ; l-HF, left heart field ; (l-)OFT, (left) outflow tract ; l-SV, left sinus venousus ; LV, left ventricle ; RV, right ventricle ; sAVC, superior atrio-ventricular canal ; ss, somite stage.

### Nodal is transiently active before heart looping, at E8.5c-e

*Nodal* expression in the lateral plate mesoderm has been previously reported at 3-6 somite stages (Vincent et al., 2004). However, we found that somitogenesis is not synchronised with heart morphogenesis (Le Garrec et al., 2017), so we re-analysed *Nodal* expression within the context of heart looping stages. By RT-qPCR of the cardiac region, we found expression of *Nodal* at E8.5c-d (Fig. 2A), which is compatible with the 3-6 somite window. We then assessed the time-window when *Nodal* is active. Because a mouse litter at E8.5 contains embryos within a range of different looping stages, we rather used drug treatment of stage-specific embryo cultures. The SB505124 drug was shown previously to efficiently abrogate the Nodal pathway (DaCosta Byfield et al., 2004; Hagos and Dougan, 2007). When applied to embryos at E8.5c over 8 hours, i.e. until E8.5e, it repressed the expression of the Nodal target *Pitx2*, whereas it only decreased it upon treatment at E8.5d and had no effect at E8.5e, compared to treatment with the adjuvant (Fig. 2B). Exposition over a shorter period of 4 hours, i.e. until E8.5d, partially reduced *Pitx2* expression. Culture of embryos over a longer period of time after treatment led to anomalies of heart looping (Fig. 2C). Taken together, these observations indicate that Nodal signalling is active between E8.5c and E8.5e, just before heart looping.

**Figure 2.**
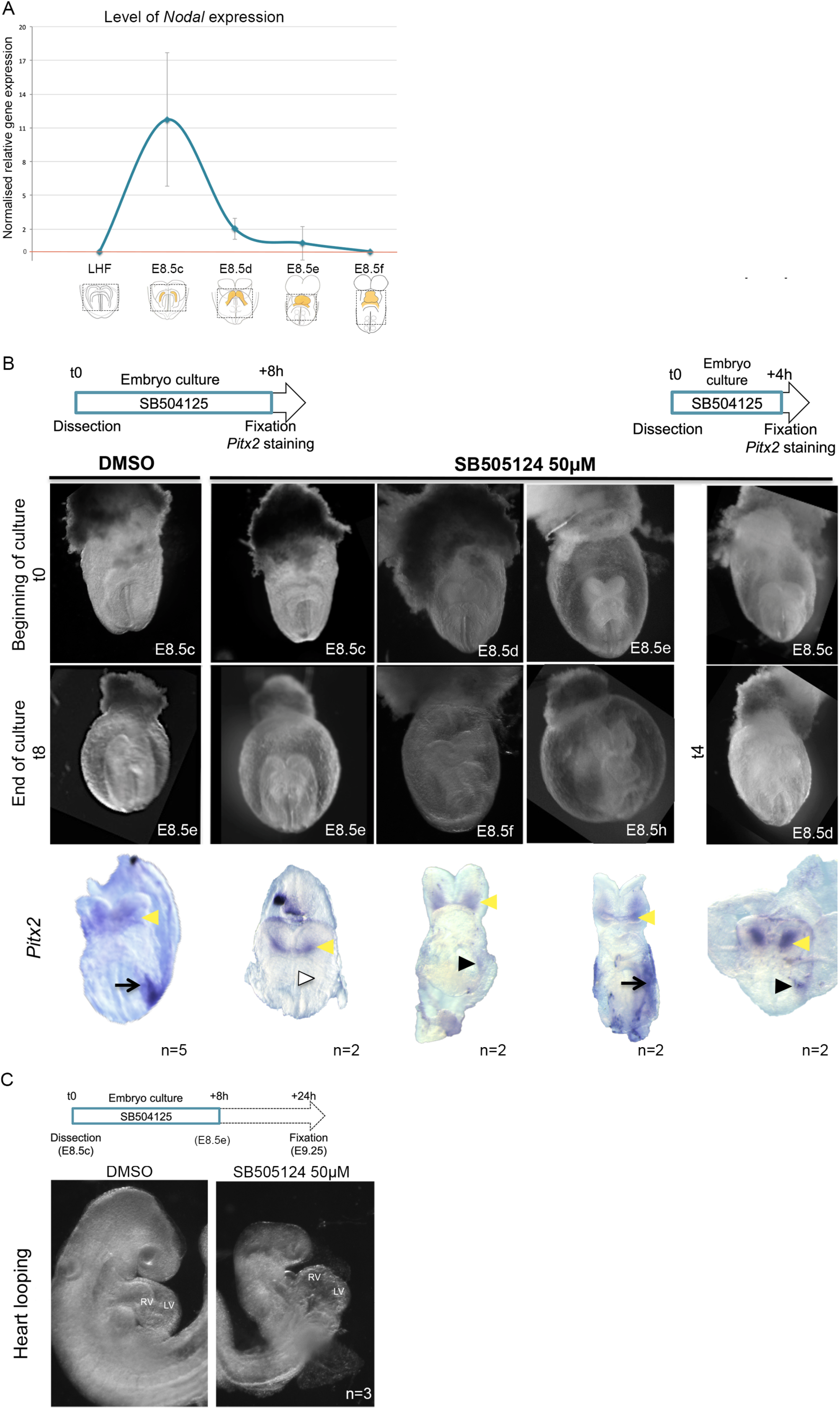
Temporal window of *Nodal* expression and signalling. (A) Quantification of *Nodal* expression by RT-qPCR in microdissected cardiac regions (dotted lines) at LHF (n=2), E8.5c (n=4), E8.5d (n=4), E8.5e (n=4) and E8.5f (n=4). The red line indicates the threshold of expression, based on primer efficiency in a reference sample. Means and standard deviations are shown. (B) The experimental design of embryo cultures is schematised, for a drug exposure of 8h (left) or 4h (right). Exemples of embryos are shown at the beginning (top panels) and end (middle panels) of the cultures, treated with the specific inhibitor of Nodal signalling (SB504125) or with the adjuvant (DMSO). *In situ* hybridisation of *Pitx2* at the end of the culture is in the lower panels, indicating strong (arrows), reduced (black arrowheads) or absent (white arrowhead) expression in the left lateral plate mesoderm, compared to control bilateral expression in the oral ectoderm (yellow arrowheads). (C) After 24h of culture, heart looping is abnormal in the presence of the drug. LHF, late headfold stage ; LV, left ventricle ; n, number of observations ; RV, right ventricle.

### *Nodal* inactivation in the mesoderm leads to four classes of looping defects

To study in more detail looping anomalies, we generated *Nodal* conditional mutants, using *Hoxb1^Cre/+^* as a driver that is expressed from the onset of gastrulation in the mesoderm (Forlani et al., 2003), overlapping with *Nodal* asymmetric expression. This provides a novel model of mesoderm-specific *Nodal* inactivation (Figure S1). We collected embryos at E9.5, when heart looping is complete in wild-type embryos, and analysed the heart phenotype of *Nodal* mutants. In a collection of 56 mutants, we observed 4 classes of anomalies (Fig. 3A), with an equal frequency (Fig. 3B). The classes are defined by the position of the ventricles : inversely lateralised (class 1), both on the right side (class 2), both on the left side (class 3) or normally lateralised (class 4). To confirm the identity of the ventricles, and more generally assess whether heart segments were correctly patterned, we performed a double *in situ* hybridisation, using *Wnt11* as a marker of the outflow tract (Zhou et al., 2007) and *Bmp2* as a marker of the atrio-ventricular canal (Ma et al., 2005) and ventral left atrium. In addition to the sulcus separating the ventricles, this was enough to identify all cardiac segments in a single labelling experiment (Fig. 3C). Staining of *Nodal* mutants indicated normal patterning of the heart tube and confirmed the localisation of the right and left ventricles in the different classes of mutants. We conclude that *Nodal* inactivation in the lateral plate mesoderm impairs the positioning of the embryonic ventricles, leading to four possible configurations, with equal frequency.

**Figure 3.**
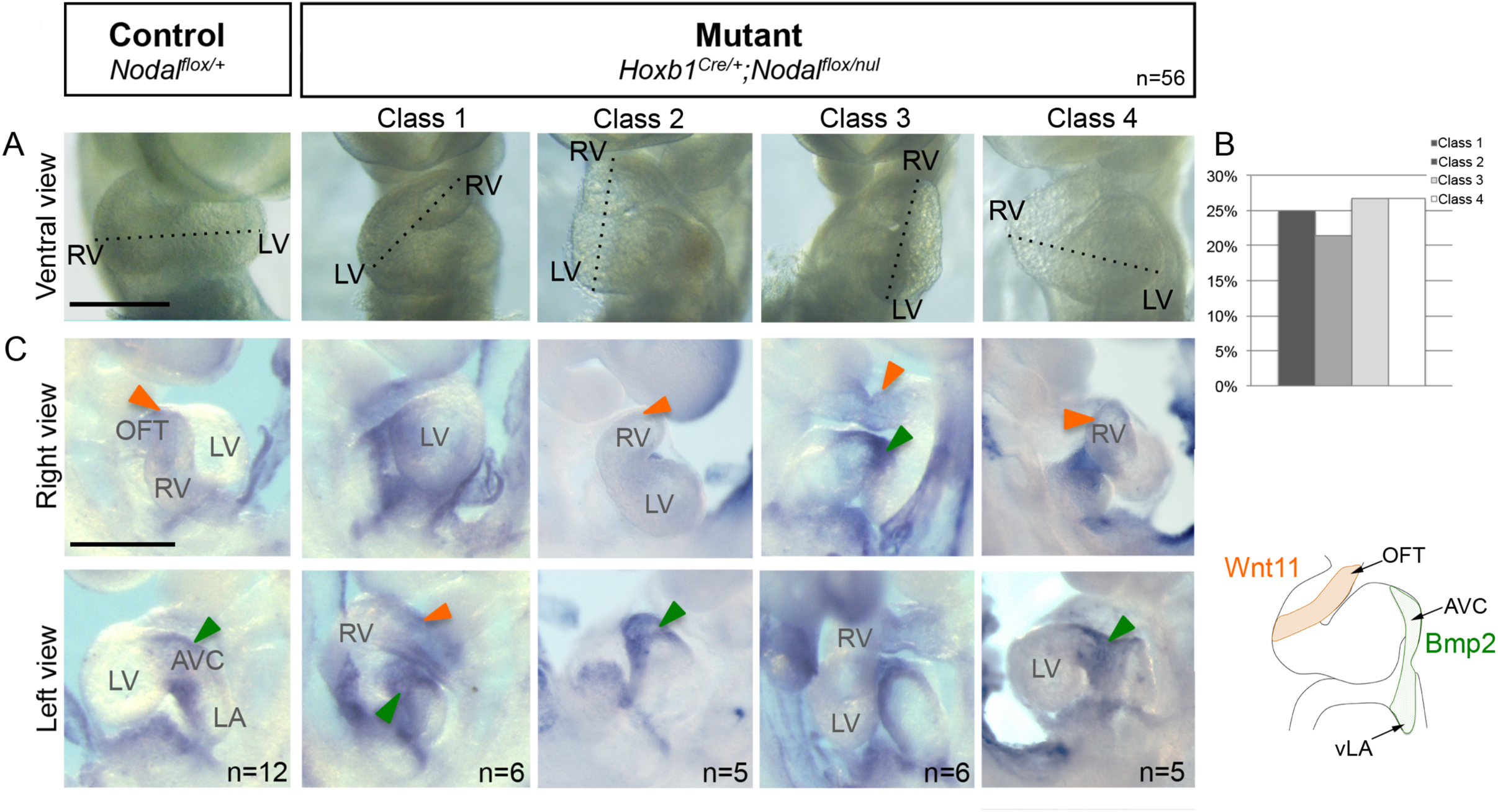
Classes of looping defects in *Nodal^flox/nul^;Hoxb1^Cre/+^*mutants. (A) Brightfield images of control and mutant embryos at E9.5. Four classes of looping defects are detected, based on the positions of the right and left ventricles. (B) Equal frequency of the four classes of mutants (p-value= 0.93, chi^2^ test, n=56 mutant embryos from 33 litters). (C) Double *in situ* hybridisation if *Wnt11* (orange arrowhead), labelling the outflow tract and the right ventricle, and *Bmp2* (green arrowhead) labelling the superior atrio-ventricular canal and the ventral left atrium. See also Movie S2 and Figure S1. AVC, atrio-ventricular canal; LA, left atrium; LV, left ventricle; OFT, outflow tract; RV, right ventricle; vLA, ventral left atrium. Scale bars: 400µm

### *Nodal* inactivation in the mesoderm randomises heart looping direction

*Nodal* inactivation was previously reported to randomise the direction of heart looping (Brennan et al., 2002; Kumar et al., 2008; Lowe et al., 2001). However, the method used to assess the loop direction was not defined. Since the looped heart tube has a helical shape, we propose to define the direction of heart looping as the orientation of the tube axis helix, seen cranially at E9.5. After HREM (High Resolution Episcopic Microscopy), 3D reconstructions of the heart loop were obtained and colour coded for cardiac segments (Fig. 4A, Movie S2). The 3D shape of the tube axis was extracted and averaged for at least 5 embryos (Fig. 4B). This shows that heart looping is rightward in class 2 and 4 *Nodal* mutants, as in control littermates, whereas it is leftward in class 1 and 3 mutants. Given the equal frequency of the mutant classes, we conclude that the direction of heart looping is indeed randomised, when *Nodal* is inactivated in the mesoderm. However, the description of the loop direction is not sufficient to characterise heart defects, since mutant classes 2 and 4 or classes 1 and 3 have distinct shapes.

**Figure 4.**
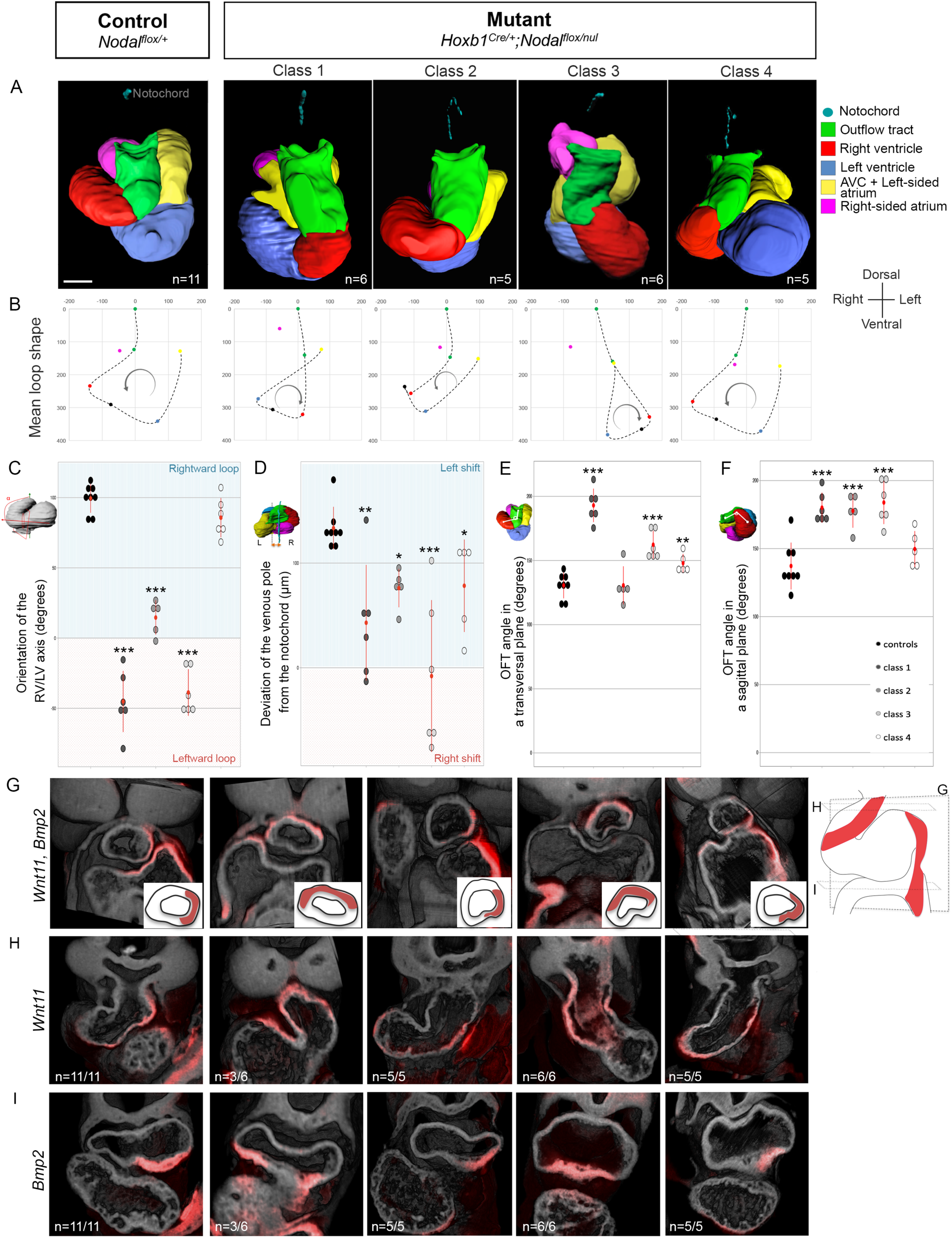
Quantification of looping defects in *Nodal^flox/null^;Hoxb1^Cre/+^*mutants. (A) Cranial views of the 3D segmented heart tube in control and mutants at E9.5, with regions colour-coded based on *Bmp2/Wnt11* expression and histology. The samples are aligned with the notochord vertical. See also Movie S2. (B) The mean trace of the tube axis is represented by a dotted line showing the shape and direction of the heart loop in controls and mutants. The origin (position 0) represents the exit of the outflow tract, seen on a transversal projection (perpendicular to the notochord). Other coloured points are the centres of gravity of the corresponding regions, with an additional point (black) at the interventricular sulcus. The arrow indicates the looping direction. (C) Quantification of the orientation of the RV/LV axis relative to the notochord in controls and the different classes of mutants. Positive (blue) and negative (orange) numbers correspond to the right and left position of the RV respectively. (D) Quantification of the lateral distance between the venous pole and the notochord. (E-F) Quantification of the curvature of the outflow tract in the transversal (E) and sagittal (F) planes. Means and standard deviations are shown. * p-value<0.05, **p<0.01, ***p<0.001 (Kruskal Wallis test, n indicated in A). See also Figure S2-S3. (G-I) 3D images by HREM of control and mutant embryos, ordered as in A. The expression of *Wnt11* and *Bmp2* is in red, the histology in grey. Section planes and the control *Wnt11/Bmp2* pattern are schematised on the right. Regionalisation of the staining in the outflow tract is schematised in the bottom right corner of upper panels. AVC, atrio-ventricular canal; L, left; LV, left ventricle; n, number of observations ; OFT, outflow tract; R, right; RV, right ventricle. Scale bar: 100µm.

### Existence of a random generator of asymmetry independent of Nodal

Within the framework of the model of heart looping that we have proposed previously (Le Garrec et al., 2017), we analysed whether the identified parameters are affected. Heart looping depends on the buckling of the heart tube, growing between fixed poles. In any class of *Nodal^flox/null^*;*Hoxb1^Cre/+^*mutants, we found no significant difference in growth compared to control littermates (Fig. S2A-F). The distance between the mutant heart poles did not significantly differ from controls (Fig. S2G). Finally, the breakdown of the dorsal mesocardium, which modulates the degree of buckling, occurred normally in the mutant samples (Fig. S2H). Thus, *Nodal* inactivation affects neither the growth nor the buckling of the heart tube.

*Nodal* mutant hearts still have an asymmetric shape, though abnormal. We quantified the degree of heart looping in *Nodal* mutants, as evidenced by the progressive repositioning of the right ventricle, from a cranial to a right location (Le Garrec et al., 2017). This measure provides a quantitative definition of the discrete classes of mutant hearts, taking into account that the angle is mirror imaged in classes 1 and 3 (Fig. 4C). In none of the mutants has this orientation remained cranio-caudal, indicating an asymmetric deformation of the tube. However, the repositioning of the right ventricle was significantly reduced in *Nodal* mutant classes 1, 2 and 3. These observations point to the persistence of some degree of heart looping in *Nodal* mutants, supporting the idea that a heart-specific random generator of asymmetry remains functional in the absence of Nodal.

### *Nodal* inactivation affects left/right asymmetries at the heart tube poles

Our previous computer simulations have shown that the buckling is biased by sequential and opposed left-right asymmetries at the arterial and venous poles. These are important to correctly shape the helix of the heart loop (Le Garrec et al., 2017). One manifestation of these asymmetries is the leftward shift of the venous pole from E8.5g. This is still detectable at E9.5 and was significantly reduced in class 2, 4 *Nodal* mutants, or abrogated (midline location) in class 1, 3 mutants (Fig. 4D). At the arterial pole, we observed anomalies in the curvature of the outflow tract at E9.5. From a cranial view, the outflow tract was significantly straighter in class 1, 3, 4 *Nodal* mutants compared to controls (Fig. 4E), whereas in a lateral view, it was straighter in class 1, 2, 3 mutants (Fig. 4F, S3A-B). Earlier at E8.5f, we detected a reduced rotation of the arterial pole in 5 mutants analysed, and at least one case of clear reversed direction (Fig. S3C).

When analysing the expression patterns of *Wnt11* and *Bmp2* in 3D, we noticed that they are regionalised in wild-type heart tube poles at E9.5, and that this is not detectable in brightfield images. *Wnt11*, which is a known target of *Pitx2* (Zhou et al., 2007), was found asymmetrically expressed on the left side of the outflow tract at E9.5 (Fig. 4G-H), in a similar domain to *Nodal-ASE-lacZ* (Fig. 1F). *Bmp2* was detected in the left, but not right, ventral atrium (Fig. 4I). So this provides molecular markers of the left/right identities of the heart poles, where sequential and opposed asymmetries are important to shape the heart loop. In *Nodal* mutants, *Bmp2* expression was found correctly left-sided in classes 2 and 4, which loop rightward. In reverse, it was abnormal in classes 1 and 3, which loop leftward (Fig. 4I). However, we observed distinct abnormal patterning of *Bmp2* in the atria : bilateral, partially penetrant, in class 1 mutants, and midline, fully penetrant, in class 3 mutants. At the arterial pole of *Nodal* mutants, *Wnt11* expression was also correctly left-sided in classes 2 and 4, whereas it was incorrectly localised to the superior outflow tract in classes 1 and 3, with a partial penetrance in class 1 (Fig. 4G-H). This indicates that the left-right patterning of the heart poles correlates with the direction of the heart loop, but is not predictive of a mutant class.

Taken together, our 3D analyses show that Nodal is required for the proper shape of the outflow tract, for the correct position of the venous pole and the right ventricle. In class1 and 3 *Nodal* mutants, leftward looping is associated with more severe anomalies and defective regionalisation of the arterial and venous poles.

### Nodal is required to amplify and coordinate left/right asymmetries

To understand the mechanism leading to the 4 classes of shapes generated in the absence of Nodal, we used our computer model of heart looping (Le Garrec et al., 2017). Based on our observations in *Nodal* mutants, we reasoned that the parameters of buckling itself were normal, whereas the left-right asymmetries at the poles were changed. Randomising the laterality of two parameters (asymmetry at the arterial or venous poles) would be sufficient to generate 4 classes of shapes. However, we also noticed that the shape of control embryos was not observed in mutants. Class 4 mutants are closest to the control phenotype, yet significantly deviate from it (Fig. 4D-E, 5A, C). In particular the right ventricle fails to reach the control right position, indicative of incomplete heart looping. In reverse, we also noticed that the leftward loops are not mirror images of the wild-type heart (Fig. 4A, 5B-C), again indicative of incomplete heart looping. Given also the reduced values of morphologial parameters (Fig. 4C-F, S3C), we postulated that left-right asymmetries at the poles not only had a variable laterality but were also reduced in intensity. Computer simulations testing these hypotheses generated the expected 4 classes of mutant shapes, as defined by the position of the ventricles (Fig. 5D compared to 3A). In addition, we quantified the orientation of the right ventricle-left ventricle axis in all classes of shapes and observed a remarkable correlation between the observed and simulated values (Fig. 5E). The model, including both a randomised laterality and a reduction of asymmetries at the poles, is thus able to recapitulate *Nodal* mutant shapes. Our work demonstrates that Nodal is required to amplify and coordinate opposed left-right asymmetries at the poles of the heart tube, thus generating a robust helical shape.

**Figure 5:**
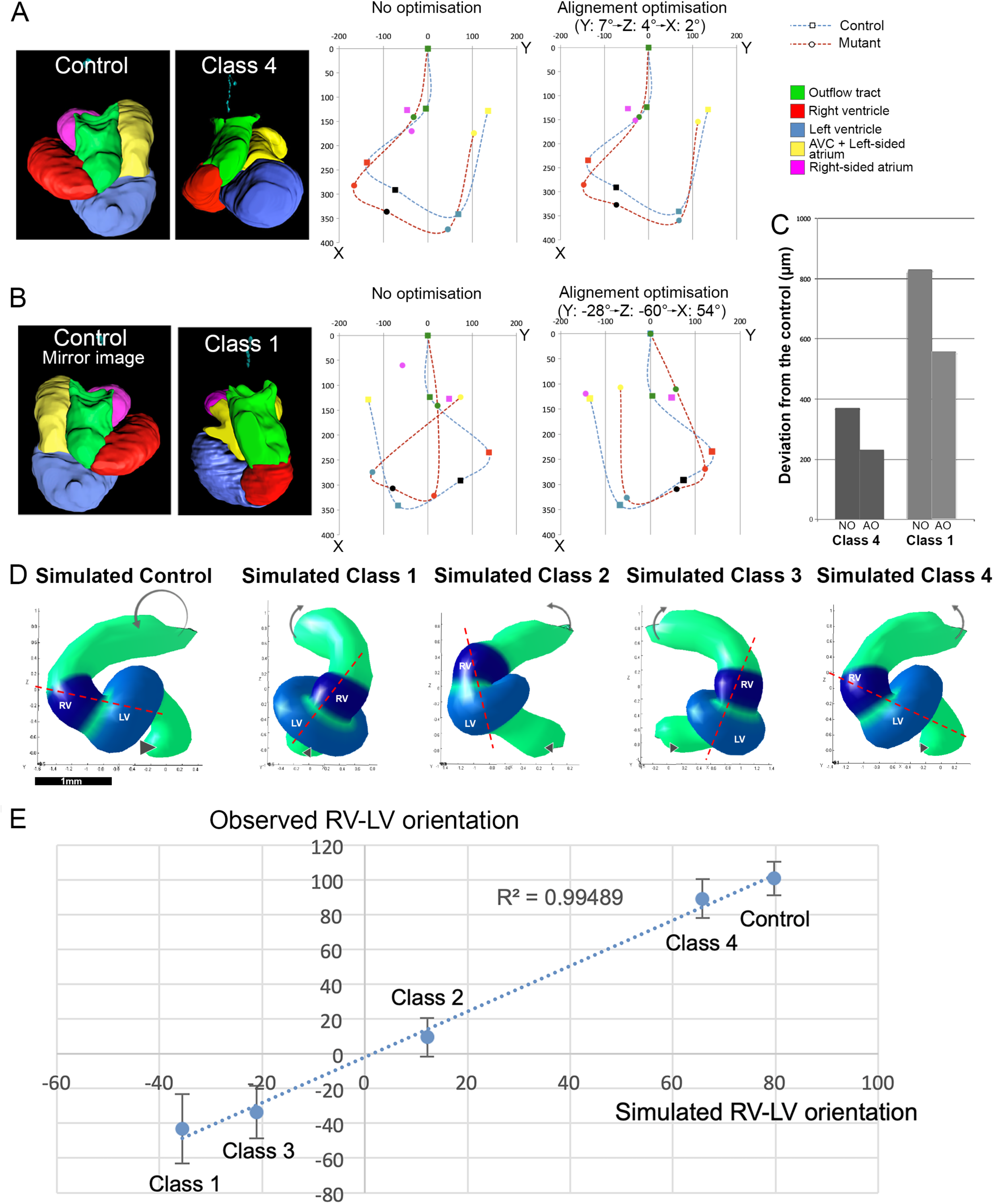
Simulations of looping defects in *Nodal* mutants by randomized laterality and reduced asymmetry at the heart tube poles. (A-C) Superimposition of the mean heart loop shape (see Fig. 4B), seen in a transversal projection, of controls and class 4 *Nodal^flox/null^;Hoxb1^Cre/+^*mutants (A), and of mirror imaged control hearts and class 1 mutants (B). The exit of the outflow tract is taken as a landmark for the alignment. The optimised alignment was obtained by sequential rotations around the three axes as indicated. (C) Quantification of the misalignment, as the total distance between the mutant and control landmark positions. (D) Computer simulations of heart looping, with different input conditions. The control simulations are run with a 25° rotation of the arterial pole (arrow) and a 2.8 fold asymmetric growth at the venous pole (arrowhead). In mutant simulations, the parameters were reduced by half. In simulated class 1, asymmetries at the arterial and venous poles were reversed, in simulated class 2 only the venous pole asymmetry was reversed, in simulated class 3 only the arterial pole asymmetry was reversed. The RV/LV axis is marked by a dotted red line, joining the centroids of the two regions. (E) Correlation (R^2^, Pearson coefficient) between the observed and simulated RV/LV axes. Means and standard deviations of the observed values are plotted (see Fig. 4C). LV, left ventricle; NO, no optimisation ; AO, alignment optimisation ; RV, right ventricle.

### Nodal modulates cell proliferation, differentiation and extra-cellular matrix composition

To understand how Nodal could modulate left-right asymmetries at a molecular level, we first analysed its main target, the transcription factor Pitx2. We quantified that *Pitx2c* is expressed as early as *Nodal* but maintained after E8.5e (Fig. S4A-B). We then analysed mutant embryos, in which the asymmetric enhancer of *Pitx2* is deleted (*Pitx2*^Δ^*ASE/*^Δ^*ASE*^^), or the asymmetrically expressed isoform *Pitx2c* inactivated. We did not detect malformations of the heart loop at E9.5, but found evidence of atrial isomerism in *Pitx2*^Δ^ASE/^Δ^*ASE*^^ hearts (Fig. S4C), as reported also downstream of *Pitx2c* (Liu et al., 2001). Surprisingly, we detected a high signal of the in situ probe against the three isoforms of *Pitx2*, suggesting potential compensation by the *Pitx2a/b* isoforms in the absence of *Pitx2c*. Thus, the *Pitx2* mutants that we have analysed do not phenocopy heart looping defects in *Nodal* mutants. To identify other targets of Nodal, we used a transcriptomic approach. The heart region (Fig. 6A) was isolated at E8.5e-f, just after *Nodal* extinction (Fig. 2A) and just before the morphological changes of heart looping. We compared the trancriptome in control and mutant samples. We first validated the dissection and genotype of samples, based on specific markers (Fig. S5A-B). Then, we analysed the GO terms of 481 differentially expressed genes (Table S1, Fig. S5C) and found an enrichment in cell cycle genes. *Ccnd2, Ccne1, Cdc25a, Cdc6, Cdt1, Mcm3/5/10, Pcna* were significantly up-regulated in *Nodal* mutants, whereas the cell cycle exit gene *Cdkn1b* was significantly down-regulated (Fig. 6B). To confirm an effect on cell proliferation, we labelled embryos with the mitotic marker phosphorylated histone H3 at different stages of heart looping (Fig. 6C). Since *Nodal* is expressed in the left heart field, we quantified mitotic cells as a ratio between the right and left heart fields. In control embryos, we observed an asymmetry in the proliferation of heart precursors, with significantly more mitotic cells on the right side from E8.5f onwards (Fig. 6D). In contrast, in *Nodal* mutants, no significant asymmetry was observed at E8.5e-i. Thus, molecular and cellular analyses support a role for Nodal in controlling the asymmetric proliferation of heart precursors. Differentially expressed genes also include genes involved in cardiomyocyte differentiation, such as *Ttn, Tnnt1* and *Vsnl1* (Fig. 6E), which are significantly down-regulated in *Nodal* mutants. More generally, in a list of 112 cardiomyocyte differentiation genes (Table S2), the vast majority is slightly down-regulated (Fig. 6E-F). With a statistical bootstrap approach, we conclude that there is a collective significant effect of *Nodal* inactivation on cardiomyocyte differentiation. As a validation, we imaged in 3D the expression pattern of *Tnnt1* and *Vsnl1*, encoding a subunit of the slow skeletal troponin and a calcium binding protein respectively (Fig. 6G-H). Expression was detected at the arterial (*Tnnt1*) and venous poles (*Tnnt1, Vsnl1). Tnnt1* was regionalized similarly to *Nodal-ASE-lacZ* (see Fig. 1F) in the left outflow tract and left atrium. Expression of both genes was decreased or bilateral in *Nodal* mutants.

**Figure 6:**
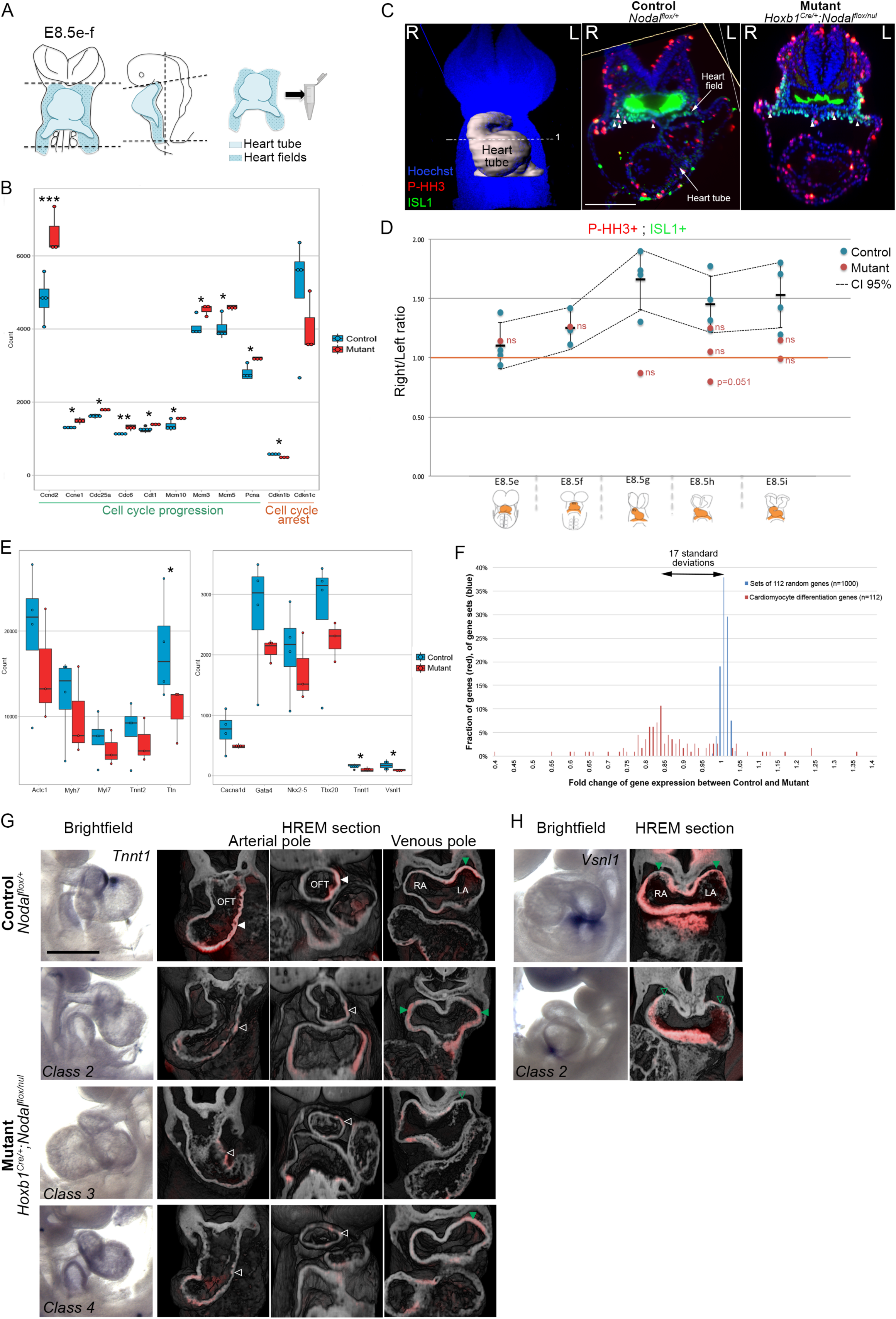
Nodal targets modulating cell proliferation and differentiation. (A) Outline (blue) of the region isolated for RNA-sequencing at E8.5e-f. (B) Normalised sequence counts of genes associated with the cell cycle, differentially expressed (fold change 1.2) in control (n=4) and *Nodal* mutant (n=3) samples. *p-value<0.05, ***p<0.001 (DESeq2). Whisker plots show the median, 25th, 75th quartiles (boxes) and the extreme datapoints (whiskers). See also Table S1. (C) Whole mount immunofluorescence of the heart field marker Isl1 and the mitotic marker Phospho-histoneH3 (P-HH3) in controls and mutants. White arrowheads point to double positive cells. L, left ; R, right. Scale bar: 200µm. (D) Quantification of cell proliferation in the right and left heart field in control (n=4, 3, 4, 4, 4 at E8.5e, f, g, h, i respectively) and mutant (n=1, 1, 1, 3, 2 at E8.5e, f, g, h, i respectively) embryos at the indicated stages of heart looping. Means, standard deviations and confidence intervals (CI) are shown for control samples. Proliferation ratios significantly deviate from 1 in control samples from the E8.5f stage, indicating proliferation asymmetry. Proliferation rates are not significantly different from a homogenous distribution between the left and right in any mutant sample (chi^2^ test of homogeneity). (E) Normalised sequence counts of genes associated with cardiomyocyte differentiation, in control (n=4) and mutant (n=3) samples. *p-value<0.05 (DESeq2). (F) Bootstrap statistical analysis to compare the fold change in the expression of 112 cardiomyocyte genes (red), between controls and mutants, with that of 1,000 randomly sampled sets of 112 genes. The mean fold change for cardiomyocyte genes (0.87) lies 17 standard deviations away from the mean of randomly sampled sets (1.01), indicating a globally significant downregulation of genes involved in cardiomyocyte differentiation. See also Table S2. (G) Brightfield images (left column) and 3D images by HREM (right columns) of *Tnnt1 in situ* hybridisation in control (upper line) and mutant (lower lines) samples at E9.5. Each line corresponds to a single embryo. The mutant class is indicated in the bottom. Filled and empty arrowheads point to high and low expression in the inner curvature of the outflow tract (white) and left atrium (green) respectively. Note that *Tnnt1* expression is bilateralised in the atria of class2 *Nodal* mutants. (H) *Vsnl1 in situ* hybridisation in control (upper line) and mutant (lower line) samples at E9.5. Filled and empty arrowheads point to high and low expression in the atria (green) respectively. LA, left atrium; OFT, outflow tract; RA, right atrium. Scale bar: 400µm. See also Figure S4-S5.

Other GO terms characterising genes differentially expressed in controls and *Nodal* mutants relate to collagen and the extra-cellular matrix (Fig. S5C). Several genes encoding extra-cellular matrix proteins, such as Col1a1/2, Col3a1, Col5a1/2, Col9a3, Reln, Tnc, the extra-cellular matrix receptor Itga2b, or the matrix metallopeptidase Mmp9, were significantly down-regulated in *Nodal* mutants (Fig. 7A). By in situ hybridisation, we found expression of *Tnc* in the heart field and the arterial pole (Fig. 7D), as previously reported (Stennard et al., 2005), whereas *Col5a2* was specifically at the venous pole (Fig. 7B), and *Mmp9,* which is required for the addition of precursor cells in the fish heart tube (Rydeen and Waxman, 2016), was detected in the mouse heart field (Fig. 7C).

**Figure 7:**
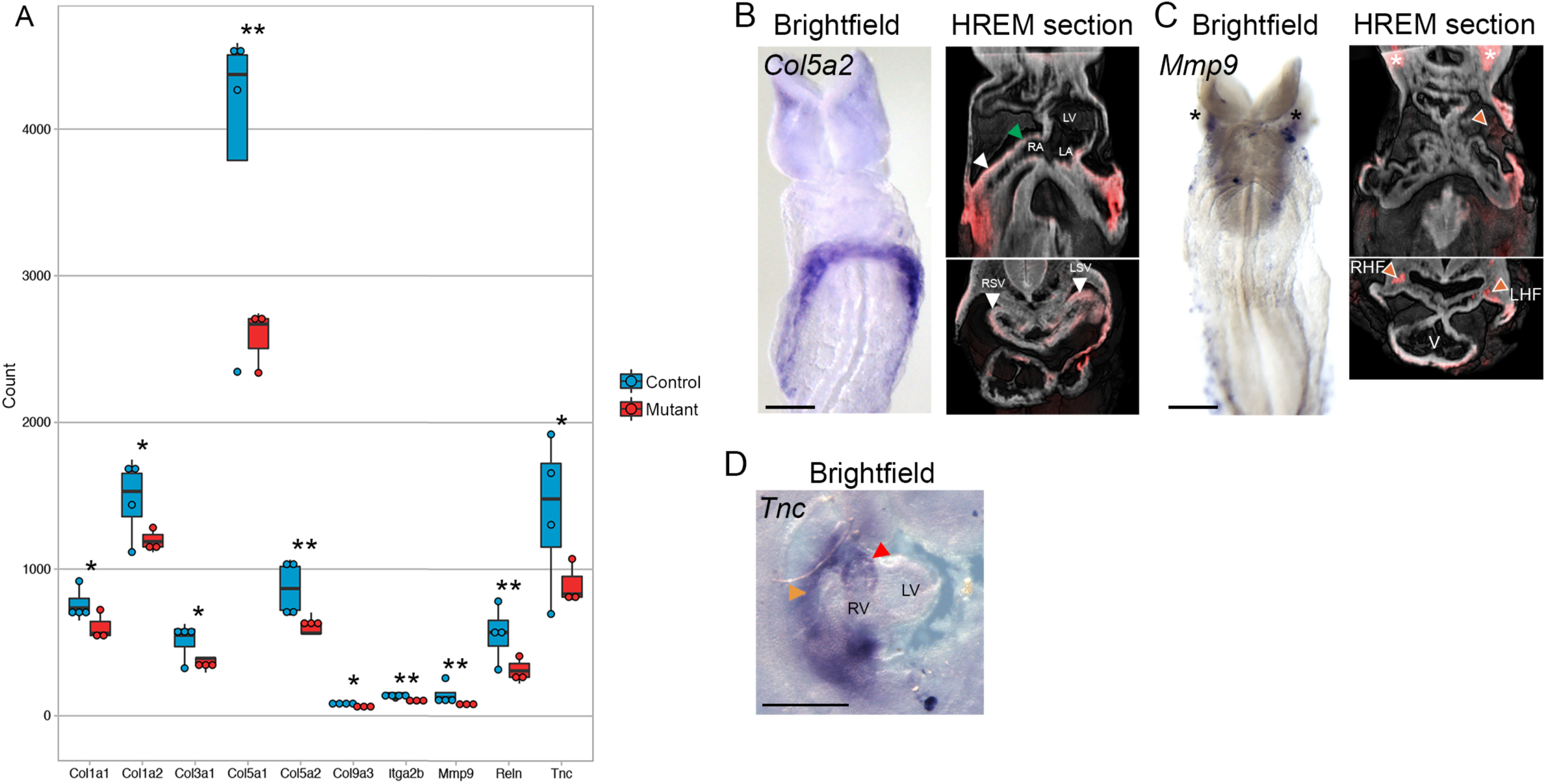
Nodal targets related to the extracellular matrix. (A) Normalised sequence counts of genes associated with the extracellular matrix, differentially expressed (fold change 1.2) in control (n=4) and mutant (n=3) samples. *p-value<0.05, **p<0.01 (DESeq2). Whisker plots show the median, 25th, 75th quartiles (boxes) and the extreme datapoints (whiskers). (B-D) Brightfield images and 3D images by HREM of *in situ* hybridisation with the indicated probes in control samples at E8.5 (B-C) or E9.5 (D). Arrowheads point to expression at the base of the atrium (green), in the sinus venosus (white), the outflow tract (red) or heart precursors (orange). Asterisks indicate labelled landmarks in the brightfield and HREM sections. LA, left atrium; LHF, left heart field ; LSV, left sinus venosus ; LV, left ventricle; OFT, outflow tract; RA, right atrium ; RHF, right heart field ; RSV, right sinus venosus, RV, right ventricle. Scale bar: 200 (B-C), 400 (D) µm.

Thus, our molecular analyses identify novel effectors of Nodal signalling, further supporting its role in amplifying left-right asymmetries at the poles of the heart tube.

## Discussion

We have mapped in 3D for the first time the contribution of *Nodal* expressing precursor cells to the heart tube poles and determined the time-window of Nodal signalling, just before heart looping. Our quantitative analyses in *Nodal* conditional mutants further demonstrate that Nodal does not initiate asymmetric morphogenesis, but rather functions as a biasing signal, to coordinate and amplify opposed left-right asymmetries at the heart tube poles. We have identified genes downstream of Nodal, which are involved in regulating local left-right asymmetries in cell proliferation, differentiation or in the composition of the extra-cellular matrix.

Nodal signalling has been initially identified as the first pathway restricted to the left side of the embryo (Collignon et al., 1996; Levin et al., 1995; Lowe et al., 1996). It is required for the asymmetric morphogenesis of several visceral organs, including the heart, lungs, spleen, stomach and gut (Brennan et al., 2002; Kumar et al., 2008; Lowe et al., 1996; Norris et al., 2002; Saijoh et al., 2003). Nodal is not only required in the node (Brennan et al., 2002), but also in the lateral plate mesoderm (Kumar et al., 2008; Saijoh et al., 2003). It is expressed transiently and turned off when the heart tube forms (Collignon et al., 1996; Lowe et al., 1996; Vincent et al., 2004). We now provide a higher spatio-temporal resolution of the expression pattern of *Nodal* in the context of heart morphogenesis, using 3D reconstructions and more resolutive staging criteria (Le Garrec et al., 2017). By stage-specific drug interference, we also show that Nodal is active during the short time window of its expression in myocardial precursor cells. *Pitx2*, a known target of Nodal, was shown to be dispensable for the proper direction of heart looping (Ammirabile et al., 2012; Liu et al., 2002; Lu et al., 1999), thus ruling out a role for mediating the biasing function of Nodal. Since, *Pitx2c* is induced as soon as Nodal is expressed we have further assessed whether *Pitx2* could be involved in another aspect of heart looping than its direction. We now show that neither the isoform *Pitx2c*, nor the *ASE* enhancer regulating asymmetric mesoderm expression, are required for the fine shaping of the heart loop, although they regulate the molecular identity of atrial chambers. Upon mutation of the three isoforms of *Pitx2* (a, b, c) heart looping is rightward, yet anomalies of the ventricles have been reported (Kitamura et al., 1999; Martin et al., 2010). Quantifications are required to investigate whether *Pitx2abc* mutants phenocopy a specific class of *Nodal* mutants. This suggests redundancy between *Pitx2* isoforms for regulating aspects of heart looping. Supporting this hypothesis, we have detected expression of *Pitx2* in *Pitx2c* mutants, as well as *Pitx2a/b* isoforms in the wild-type heart region at E8.5e/f (RNA-seq data, not shown). Expression of *Pitx2a/b* had not been detected previously by in situ hybridisation (Kitamura et al., 1999; Liu et al., 2001; Schweickert et al., 2000), probably reflecting a less sensitive approach and/or less resolutive staging. Whether Pitx2 mediates aspects of Nodal role for heart looping thus remains an open question. Our transcriptomic analysis identifies novel targets of Nodal signalling, regulating the proliferation, differentiation and extra-cellular matrix composition of cardiac cells. Several of these targets (*Tnnt1*, *Vsnl1*, *Col5a2* or *Tnc*) are expressed in the heart tube poles or heart field, in agreement with the contribution of *Nodal* expressing cells. Our transcriptomic data highlight downregulation of many genes involved in myocardial differentiation, including some previously identified in the context of left-right asymmetry, such as *Acta1* (Noël et al., 2013) or *Hcn4* (Pai et al., 2017). Previous targets of Nodal signalling had been identified in ES cells or during fish gastrulation and mesendoderm specification (Bennett et al., 2007; Brown et al., 2011; Coda et al., 2017; Guzman-Ayala et al., 2009). We have detected little overlap with our gene list (Nodal pathway components, *Cyp26a1*), in keeping with distinct roles of Nodal in distinct tissues and at distinct stages. A common theme is a role of Nodal in cell differentiation, and potentially in cell proliferation and re-arrangements, relevant also to Nodal re-expression during cancer progression and metastasis (see Quail et al., 2013).

The phenotype of asymmetry can be decomposed into several components: the initiation of the asymmetry, the fold difference between the left and right, and the laterality of the difference, i.e. whether a left determinant is localized on the anatomical right or left side. Our quantitative analysis of heart looping in *Nodal* mutants, combined with computer simulations, demonstrates that Nodal is not required to initiate asymmetric heart looping, and that it rather functions as a biasing signal, to coordinate the laterality of asymmetries at the tube poles and thus generate a robust helical shape. In the absence of *Nodal*, heart looping is not randomized in the sense that specific shapes are generated, rather than a continuum of random shapes. The observation of 4 classes of heart loop shapes with an equal frequency supports the model that independent left-right asymmetries at the two heart poles determine the loop shape. It is the laterality of these asymmetries which is randomised in *Nodal* mutants, but not the process of asymmetry *per se*. Such a biasing role for Nodal has also been reported for the stomach (Kumar et al., 2008; Saijoh et al., 2003), or for the migration of the parapineal nucleus in the fish diencephalon (Concha et al., 2000). In other instances, Nodal is required to initiate asymmetry and its absence leads to symmetrical phenotypes or isomerism. This is the case for the formation of the mouse lungs and spleen (Brennan et al., 2002; Kumar et al., 2008; Lowe et al., 2001; Saijoh et al., 2003), or for the expression of *cxcr4b*, an early marker of the left habenular nucleus in the fish diencephalon (Roussigne et al., 2009). Our quantitative analyses of shape further show that *Nodal* mutant hearts are distinct from the control shape, when left-right asymmetries are correctly lateralised (Class 4), and that *Nodal* mutant hearts are also distinct from a mirror image, when left-right asymmetries are inversely lateralised (Class 1). This demonstrates a novel role for Nodal in amplifying left-right asymmetries. Our identification of genes downstream of Nodal and involved in cell proliferation, differentiation and extra-cellular matrix composition at the heart tube poles, provides molecular candidates for mediating this amplification. The fact that the reversal of asymmetries does not generate a perfect mirror image of organs may explain why situs inversus totalis (incidence 3/100,000) is not always asymptomatic, but can be associated with anomalies in the heart, spleen and intestinal rotation (Lin et al., 2014).

Our study supports the two-step model of asymmetric organogenesis proposed by Brown and Wolpert (1990). In the absence of *Nodal*, we show that some degree of asymmetric morphogenesis occurs, as reported also upon bilateral expression of *Nodal*, in *Snai1* mutants (Murray and Gridley, 2006). This is in agreement with the existence of a generator of asymmetry that is independent of the Nodal left-right bias. We had shown previously that heart looping can be reproduced by a buckling mechanism, when the heart tube grows between fixed poles (Le Garrec et al., 2017). Our computer simulations have shown that this mechanism in isolation generates random deformations, whereas small left-right asymmetries at the tube poles are sufficient to bias them. We thus provide the first demonstration of the concept of an organ-specific random generator of asymmetry, with a simple buckling model able to generate a range of shapes from a limited number of initial parameters. Whether this model also applies to other asymmetric tubular organs such as the gut remains to be investigated. Another pending question is the mechanism of the generator of asymmetry. Our work points to the existence of left-right asymmetries at the heart tube poles, independent of Nodal, which orient the buckling of the heart tube. We have previously characterised a rightward rotation of the arterial pole and asymmetric cell ingression at the venous pole (Le Garrec et al., 2017). We now show that some rotation of the arterial pole is still detectable in the absence of Nodal, with a leftward or rightward laterality. However, the underlying molecular mechanism remains unknown. The asymmetric cell proliferation that we have observed in the heart field from E8.5g depends on Nodal signalling and thus would not fit as a mechanism. Actomyosin, independently of Nodal, was shown to be important for the later phase of heart looping in the fish (Noël et al., 2013). By homology with other rotation processes, one can speculate about other mechanisms. Directional movements of myocardial cells are involved in the rotation of the zebrafish cardiac cone (Baker et al., 2008; Rohr et al., 2008; Smith et al., 2008), whereas rotation of the fly hindgut is associated with the chiral activity of the atypical myosin Myo1d, propagated by Planar Cell Polarity (González-Morales et al., 2015). Asymmetries preceding that of Nodal are evident in the brain, involving Fgf, Notch or Wnt/β-catenin signalling (see Güntürkün and Ocklenburg, 2017), or in the frog embryo, involving ion channels or cytoskeletal rearrangements (Pai et al., 2017).

In the context of heart morphogenesis, we now unravel the specific contribution of Nodal to heart looping. We show that *Nodal* is transiently expressed in cardiac precursor cells and required during this time window, E8.5c-e, for heart looping. *Nodal* expressing cells contribute to the myocardium of the arterial and venous poles, but neither to the left ventricle, nor to the majority of the right ventricle. This is reminiscent of the second myocardial lineage (Meilhac et al., 2004) and is consistent with fate mapping of the left second heart field (Domínguez et al., 2012). We now provide a 3D map of left derivatives in the looped heart tube. Given the short time window of *Nodal* expression and the perdurance of β-galactosidase a day later, we expect that most of *Nodal* expressing cells in the lateral plate mesoderm are labeled with the *Nodal-ASE-lacZ* transgene, compared to an hypothetical *Nodal-ASE-Cre* transgene, which would induce a delay in the initial activation of a reporter. Our observations of largely negative ventricles would thus suggest that cells of the first lineage are not sensitive to left-right patterning. They are defined first (Meilhac, Dev Cell 2004), ingress first in the primitive streak (Lescroart et al., 2014), differentiate earlier (Ivanovitch et al., 2017) and reach the heart region (cardiac crescent) before the node has become a left-right organizer at the LHF stage (Kawasumi et al., 2011) and before the heart field expresses *Nodal* at E8.5c. Our quantification of the coverage of the *Nodal-ASE-lacZ* staining suggests a 25% contribution of *Nodal* expressing cells to the heart poles at E9.5. This percentage below 50% raises the questions of whether all left cells in the second heart field express *Nodal*, and/or whether left precursor cells contribute less to the heart tube compared to right cells. The higher proliferation of cells that we detect in the right second heart field, as well as the asymmetric contribution of right versus left grafts in the chick precardiac mesoderm (Stalsberg, 1969) would support the second hypothesis. The fact that *Nodal* is expressed in heart precursors and active before heart looping supports the idea of an extrinsic role of Nodal for heart looping, i.e. in cells outside the heart tube. In contrast its target *Pitx2c* is expressed within the heart tube, in an asymmetric domain largely overlapping with that of *Nodal-ASE-lacZ*, with the exception of the ventricles, in which more cells positive for *Pitx2c-ASE-lacZ* are detectable basally (Furtado et al., 2011; Shiratori et al., 2006). *Pitx2c* was thus thought as a good candidate to mediate an intrinsic role of Nodal signalling. However, its contribution to heart looping remains unclear and more generally whether there are other intrinsic determinants of heart looping remains to be demonstrated. We provide here and previously (Le Garrec et al., 2017) many quantifiable parameters of heart looping, which will be important to compare anomalies in different experimental conditions.

Our work provides novel insight into the left-right patterning of cardiac cells, showing that it is a spatio-temporal dynamic process rather than a single event. We have previously detected opposed and sequential left-right asymmetries at the arterial and venous poles (Le Garrec et al., 2017). We now report molecular left-right patterning of the outflow tract (*Wnt11*) and atria (*Bmp2*), which is defective in *Nodal* and *Pitx2^Δ^ASE^^* mutants. However, anomalies in molecular patterning are not fully penetrant and not predictive of a class of abnormal heart looping. This raises the possibility of other levels of left-right asymmetry in the heart, which are regulated independently, given the absence of strict correlations. With more markers of the heart tube poles identified in our transcriptomic analysis, the molecular signature of the asymmetry of cardiac segments will be refined. Another cardiac asymmetry is the curvature of the outflow tract, arising between E8.5j and E9.5, under the control of Nodal. This curvature is not reproduced in our computer simulations, suggesting that it is not related to the buckling. It is also not correlated with proliferation differences (data not shown), leaving its origin unclear.

In the clinics, heterotaxy is a laterality defect associated with mutations affecting the formation or signalling of the left-right organizer (Guimier et al., 2015). The clinical picture is very variable, in terms of associations of left-right anomalies between different organs or between different heart segments (Desgrange et al., 2019; Lin et al., 2014). Even looking at a single structure such as the atria, the parameters of the anatomy of the appendages and of the connection of the inferior caval vein are not always concordant (Tremblay et al., 2017). This is the basis of debates in the nomenclature of heterotaxy, for the qualification of isomerism (Jacobs et al., 2007). Clinical variability may relate to the fact that left-right asymmetry is regulated independently at different levels, and that Nodal regulate either the initiation or the laterality of asymmetry. The previous focus on the symmetry-breaking event has masked the dynamics of left-right patterning. Quantifications of the contribution of different factors to the fine shape of the heart loop, not just its direction, are expected to provide novel perspectives in understanding the origin of severe congenital heart defects associated with heterotaxy or other types of cardiac chamber misalignment.

## Author Contributions

Conceptualization: AD, JFLG, SMM; Methodology: AD, JFLG; Software : JFLG ; Formal Analysis : JFLG, AD ; Investigation: AD, JFLG, SB, TB; Writing - original draft: SMM; Writing - review & editing: AD, JFLG, SMM; Visualization: AD, JFLG, SB ; Supervision: SMM, AD ; Project administration: SMM ; Funding acquisition: SMM.

## Acknowledgments

We thank V. Benhamo, L. Guillemot, L. Bombardelli, M. Bertrand, A. Murukutla, V. Nikovics and C. Cimper for technical assistance, JN. Domínguez for advice in embryo culture, J. Collignon for *Nodal-ASE-lacZ* embryos, H. Hamada and Y. Ikawa for *Pitx2*^Δ^*ASE/*^Δ^*ASE*^^ embryos, N. Kurpios for *Pitx2c^null/+^* mice, L. Robertson for *Nodal^flox/flox^* mice, S. Zaffran for *Hoxb1^Cre/+^* mice, the histology and imaging platforms of the SFR Necker, C. Bole-Feysot and O. Alibeu of the Genomics platform, N. Cagnard of the Bioinformatics platform, C. Dicu and the LEAT animal facility. This work was supported by core funding from the Institut *Imagine* and Institut Pasteur, a grant from the ANR [16-CE17-0006-01] to SMM and the MSD-Avenir fund (Devo-Decode). SB has benefited from a fellowship of the Société Française de Pédiatrie, TB was supported by the Pasteur - Paris University International PhD Program, SMM is an INSERM research scientist.

## Material and Methods

### EXPERIMENTAL MODEL AND SUBJECT DETAILS

#### Animals

Wild-type mouse embryos were from a C57Bl6J genetic background. The *Nodal-ASE-lacZ* transgenic line (Norris and Robertson, 1999), *Nodal^flox/flox^* (Lu and Robertson, 2004), *Pitx2*^Δ^*ASE/*^Δ^*ASE*^^ (Shiratori et al., 2006), *Pitx2c^null/null^* (Liu et al., 2002) lines were maintained in a mixed genetic background. *Nodal^null/+^* mice were generated by crossing *Nodal^flox/flox^* males with *Mef2cAHFCre* transgenic females (Verzi et al., 2005) and then crossed to *Hoxb1^Cre/+^* (Arenkiel et al., 2003). *Nodal^null/+^;Hoxb1^Cre/+^* males were maintained in a mixed genetic background and crossed to Nodal*^flox/flox^*females to generate *Nodal* conditional mutants. Both male and female embryos were collected and used randomly for experiments. Embryonic day (E) 0.5 was defined as noon on the day of vaginal plug detection. Heart looping stages from E8.5c to E8.5j were defined according to the previously published nomenclature (Le Garrec et al., 2017), whereas E8.5a and E8.5b are equivalent to EHF and LHF stages respectively (Downs and Davies, 1993). The number of somites was evaluated from the HREM images. All embryos were genotyped by PCR. For the genotyping of Nodal alleles, the reverse primer CCTGACTCAAAACCCAAGGC was used in combination with the forward primer ATTCCAGCAGTTGAGGCAGA to detect the wild-type (540b) or floxed (590b) alleles and the forward primer CCACCCAATTTCTAGCCCAG to detect the deleted allele (1050b). Animal procedures were approved by the ethical committees of the Institut Pasteur and Paris Descartes and the French Ministry of Research.

### METHOD DETAILS

#### Embryo culture

For drug treatment, wild-type E8.5 embryos were collected in Hank’s solution. We tested a range of drug concentrations (20-50µM) to avoid toxicity on embryo development and promote efficient *Pitx2* downregualtion. A working concentration of 50µM of SB505124 or an equivalent volume of the adjuvant (DMSO) were added to the 75% rat serum, 25% T6 medium, supplemented with 1X Penicillin/Streptomycin. Embryos were cultured with 5% CO2, 5% O2, in rolling bottles in a precision incubator (BTC Engineering, Milton, Cambridge, UK). At the end of the treatment, embryos were rinsed in PBS and fixed in paraformaldehyde (PFA) 4% or further incubated in fresh culture medium, in a 5% CO2-20% O2 atmosphere, and harvested 24h after the initiation of the culture. Brightfield images were acquired at the beginning and the end of the culture with a Zeiss AxioCamICc5 Camera and a Zeiss StereoDiscovery V20 stereomicroscope with a Plan Apo 1.0X objective.

#### Wholemount β-galactosidase staining and immunofluorescence

Embryos were collected at E8.5 or E9.5. The heart was arrested in diastole with 250mM KCL (E9.5). *Nodal-ASE-LacZ* transgenic embryos were fixed in 4% PFA – 5mM EGTA – 2mM MgCl2 for 10min. Embryos where then permeabilized in 0.2% NP40 – 2mM MgCl2 – 0.1% sodium deoxycholate 30min and stained overnight in Xgal solution. Immunofluorescence on whole mount E8.5 embryos was performed after removal of the left headfold as a landmark, using CUBIC clearing as described in (Le Garrec et al., 2017). Multi-channel 16-bit images were acquired with a Z.1 lightsheet microscope (Zeiss) and a 20X/1.0 objective. Automatic detection of mitotic cells was performed with the Spots plugin of Imaris and co-localisation with Isl1 staining was evaluated manually. Given morphological variations in mutant embryos, their stage of development was evaluated based on the length of the heart tube, using the quantifications of (Le Garrec et al., 2017) as a reference. P-Smad2 staining was adapted from Kawasumi et al, 2011. E8.5 embryos were fixed 2h in 4% PFA, and gradually dehydrated into methanol. We used 3% H2O2 for bleaching before rehydratation. Samples were blocked with the TSA Blocking Reagent (Perkin Elmer), incubated with the primary antibody Phospho-Smad2 (1/50) during 48h at 4°C and 4h with the Alexa Fluor conjugated secondary antibody (1/500). Embryos were then transparised in gradually concentrated glycerol before imaging with a fluorescent Stereomicroscope in 80% gycerol.

#### RT-qPCR

Cardiogenic regions were isolated either as shown in Fig. 6A or by removing the heart tube (from E8.5e) and cutting the heart field to keep the left halve only. The posterior boundary is set at the level of the second somite, in agreement with fate maps (Domínguez et al., 2012). The tissue was flash frozen in liquid nitrogen. RNAs were extracted in TRIzol-Chloroform and purified using the RNeasy micro kit. Reverse transcription was carried out using the Quantitect Reverse Transcription kit. Quantitative PCR was carried out using a real-time PCR system (BioRad), and primers as listed in the Key Resources Table. *Polr2b* was chosen from the RNA-seq dataset as a reference housekeeping gene, because of its expression in the range of Nodal pathway components (2000 counts), with no variability between samples, including controls and mutants. The mRNA expression levels were measured relatively to *Polr2b* and normalized with a reference cDNA sample (pool of 4 embryos at E8.5c d, g, and j), using the standard ΔΔCt method.

#### RNA in situ hybridisation

*In situ* hybridisation was performed on wholemount embryos after fixation in PFA 4% and dehydratation in methanol 100% following standard protocols. *Lefty2, Pitx2, Wnt11, Bmp2, Sema3c*, *Mmp9*, *Vsnl1* riboprobes were transcribed from plasmids. *Col5a2, Tnc* and *Tnnt1* probes were synthetized by PCR, using primers listed in the Key Resources Table. Hybridization signals were detected by alkaline phosphatase (AP)-conjugated anti-DIG antibodies (1/2500; Roche), which were revealed with NBT/BCIP (magenta) substrate (Roche). After staining, the samples were washed in PBS and post-fixed.

#### HREM (High-Resolution Episcopic Microscopy)

Embryos were collected and embedded, as described in (Le Garrec et al., 2017), in methacrylate resin (JB4) containing eosin and acridine orange as contrast agents. Two channels images of the surface of the resin block were acquired using the optical high-resolution episcopic microscope (Indigo Scientific) and a 1X Apo objective repeatedly after removal of 1.56-1.7 µm thick sections : the tissue architecture was imaged with a GFP filter and the staining of enzymatic precipitates with a RFP filter. The dataset comprises 500-1700 images of 1.15-1.85 µm resolution in x and y depending on the stage. Icy (de Chaumont et al., 2012) and Fiji (ImageJ) softwares were used to crop or scale the datasets. 3D reconstructions were performed with the Fiji plugin Volume Viewer or Imaris (Bitplane).

#### RNA sequencing

Embryos were microdissected and the tissue was flash frozen in liquid nitrogen. RNA was extracted in TRIzol-Chloroform and purified using the RNeasy micro kit. RNA quality and quantity were assessed using RNA Screen Tape 6000 Pico LabChips with the Tape Station (Agilent Technologies). All RINs were higher than 9.0. The library was established using the Nugen Universal Plus mRNA-Seq kit, using 20 ng of total RNAs as recommended by the manufacturer. The oriented cDNAs produced from the poly-A+ fraction were PCR amplified (15-18 cycles). An equimolar pool of the final indexed RNA-Seq libraries was sequenced on an Illumina HiSeq2500, with paired-end reads of 130 bases and a mean sequencing depth of 58 millions per sample. The RNA-seq data have been submitted to the NCBI Gene Expression Omnibus (GEO).

#### FEA Modeling of the heart tube

The model is based on the GFtbox finite element analysis software, using a cylindrical mesh, as described in (Le Garrec et al., 2017), with fixed poles and a dorsal constraint simulating the progressive breakdown of the dorsal mesocardium. At each successive step during a simulation, each element is deformed according to a growth tensor field specified from the hypotheses of the model.

The control model is based on the following input parameters : basic longitudinal growth (2.5%, with a peak value of 5% ventrally between steps 1-40), 25 degrees rightward rotation at the arterial pole (1.1% circumferential growth per step, positive on the left side, negative on the right side), asymmetric longitudinal growth at the venous pole (peak value of 7% on the right, 2.5% on the left), circumferential growth in the ventricles (0.9% in the ventral right ventricle, 0.4% in the dorsal right ventricle, 1.4% in the left ventricle). Simulations were run for 100 steps. The MATLAB code containing the interaction function of the GFtbox model, and used to generate the control shape in Fig. 6A, is provided in *Source code 1*.

The simulations of the mutant shapes were obtained by a 50% reduction in the intensity of the asymmetries at both poles and by simulating the four possible combinations of lateralization at the arterial/venous poles : normal/normal, normal/inverted, inverted/normal and inverted/inverted. The MATLAB code containing the interaction function of the GFtbox model, and used to generate the mutant shapes in Fig. 6A is provided in *Source code 2*. This code is edited to run the inverted/inverted combination (“Class 1”), the alternative codes for each of the three other combinations being included as commented lines under the headings “Class 2-4”.

### QUANTIFICATION AND STATISTICAL ANALYSIS

#### Quantification of the proportion of β-galactosidase staining

Hearts were segmented from HREM images using the IMARIS software (Bitplane). The contour of the myocardium was manually outlined at regular Z intervals of the GFP channel, and the Create Surface function was used to reconstruct the 3D surface. Signal of the β-galactosidase staining intersecting with the myocardium was obtained using the “mask selection” function to extract a new channel and create another 3D surface. The volume of each surface was extracted and the proportion of the stained myocardium was calculated.

#### 3D rendering and visualisation

We used Fiji Volume Viewer plugin to assess in situ labeling within the heart tube in 3D. In parallel we segmented the myocardium from HREM using the IMARIS software (Bitplane). This 3D surface was used to extract the RFP channel corresponding to in situ signal within the object. The extracted signal was used to automatically generate a corresponding 3D surface.

#### Quantification of the geometry of the heart loop

From the 3D reconstruction of the myocardium contour, the axis of the cardiac tube was extracted, using eight landmarks along the length of the tube. Three of these landmarks were obtained with the IMARIS Oblique Slicer function intersecting the tube perpendicularly, and by computation of the centroid of the polygon (MATLAB geom3d library : function polygonCentroid3d) : one at the exit of the outflow tract, one at the sulcus between the two ventricles, and one at the bifurcation of the two atria. The five other landmarks were obtained by sub-division of the volume into the outflow tract (positive for *Wnt11*), the right ventricle (without cushions), the left ventricle, the atrio-ventricular canal and left atrium (positive for *Bmp2*) and the right atrium. The center of gravity of each of these volumes was computed, using IMARIS.

The 3D coordinates of the eight landmarks were used to draw the loop of the tube axis as shown in Fig. 4. All the hearts were aligned so that the notochord coincides with the Z axis, and the dorsal-ventral axis with the X axis. This alignment was performed using two landmarks on the notochord and two landmarks defining the bisectrix of the neural groove, then applying two successive 3D rotations to the axis coordinates, using an in-house MATLAB code: first a rotation aligning the notochord with the Z axis, then a rotation aligning the dorsal-ventral axis with the X axis. All measurements shown in Fig. 4 were done on these alignments and averaged for all analysed samples (n indicated in Fig. 4A). The orientation of the RV/LV axis relative to the notochord, the distance of the venous pole (taken as the bifurcation of the tube) relative to the notochord, the distance between the poles and the tube length, the rotation of the tube at E8.5f were calculated as in Le Garrec et al., 2017. The OFT angles were directly measured on the aligned loops after projection on the indicated planes (transversal : XY; sagittal : XZ).

Superimposition of the loops shown in Fig. 5 was obtained by optimization of composite 3D rotations (combining rotations around the three coordinate axes) of the mutant loops relative to the control loop. The 3 basic rotation matrices were each successively incremented by 1° and combined in alternative order (because of non-commutation) to explore the full space of 3D rotations around the exit of the outflow tract taken as a fixed point. Optimization was obtained by minimizing the sum of the euclidean distances between each of the seven mutant landmarks and the corresponding control landmarks. The computations were implemented in MATLAB.

#### Bioinformatics analyses of the RNA sequences

FASTQ files were mapped to the ENSEMBL [Mouse GRCm38] reference using Hisat2 and counted by featureCounts from the Subread R package. Read count normalisations and group comparisons were performed by three independent and complementary statistical methods : DESeq2, edgeR and LimmaVoom. Flags were computed from counts normalized to the mean coverage. All normalized counts <20 were considered as background (flag 0) and >=20 as signal (flag=1). P50 lists used for the statistical analysis regroup the genes showing flag=1 for at least half of the compared samples. Unsupervised cluster analysis was performed by hierarchical clustering using the Spearman correlation similarity measure and average linkage algorithm. The results of the three methods were filtered for differentially expressed genes between control and mutant samples, on the basis of a p-value lower than 0.05 and a fold change greater than 1.2. Functional analyses were carried out using the Gene Ontology database (PANTHER Overrepresentation test).

#### Bootstrap inference on transcriptomic data

In order to assess how significantly genes of cardiomyocyte differentiation deviate from the average gene expression fold change between control and mutant samples, a bootstrap method was applied. The analysis was restricted to the 9,375 genes with a minimum of 150 normalised counts (see Fig. S5B). 112 were selected as cardiomyocyte differentiation, because they encode sarcomere components, ion channels, transcription factors, adhesion proteins or signals required for cardiomyocyte differentiation, or on the basis of previous RNA-seq datasets (DeLaughter et al., 2016; Li et al., 2016). The 9,375 genes were resampled 1,000 times, with replacement, into sub-samples of 112 genes (MATLAB RandStream method using the Mersenne twister generator). The means and standard deviations of the fold change distribution between mutant and control embryos for these 1,000 bootstrap samples were assessed, and compared to the mean and standard deviation for the 112 genes of cardiomyocyte differentiation.

#### Statistical analysis

Sample size was checked post-hoc, using the calculator powerandsamplesize.com, in order to ensure a power of at least 0.8, with a type I error probability of 0.05, with an effect size of 20%. The collection of full litters was used to randomise imaging experiments. Group allocation was based on PCR genotyping. 3 outliers were excluded from geometric analysis based on a lower number of somites (Fig. 4 and S2). One mutant sample was discarded from the RNA-seq analysis because of a poor gene coverage. All sample numbers (n) indicated in the text refer to biological replicates, i.e. different embryos. Investigators were blinded to allocation during imaging and phenotypic analysis, but not during quantifications. Tests were performed with Excel and R. The correlation between two data series was quantified by the square of the Pearson coefficient R^2^. The regression line was computed using the least square method. Comparisons of two centre-values were done on the average, or the geometrical mean when ratios were compared, using a Student two-tailed test. When more than two centre-values were compared, an ANOVA was calculated, unless a normal distribution could not be assumed, in which case a Kruskal Wallis test was used. For comparing left and right angles at successive positions, a paired Student test was used. The 95% confidence intervals for the mean were calculated assuming a normal distribution of measurements. A chi^2^ test was used to evaluate the randomisation of class frequency and the symmetry of proliferation rates between left and right heart precursors.

